# Expansion and diversification of the gibberellin receptor GIBBERELLIN INSENSITIVE DWARF1 (GID1) family in land plants

**DOI:** 10.1101/221937

**Authors:** Rajesh K. Gazara, Kanhu C. Moharana, Daniel Bellieny-Rabelo, Thiago M. Venancio

## Abstract

Gibberellic acid (gibberellins, GA) controls key developmental processes in the life-cycle of land plants. By interacting with the GIBBERELLIN INSENSITIVE DWARF1 (GID1) receptor, GA regulates the expression of a wide range of genes through different pathways. Here we report the systematic identification and classification of GID1s in 52 plants genomes, encompassing from bryophytes and lycophytes, to several monocots and eudicots. We investigated the evolutionary relationship of GID1s using a comparative genomics framework and found strong support a previously proposed phylogenetic classification of this family in land plants. We identified lineage-specific expansions of particular subfamilies (i.e. GID1ac and GID1b) in different eudicot lineages (e.g. GID1b in legumes). Further, we found both, shared and divergent structural features between GID1ac and GID1b subgroups in eudicots, which provide mechanistic insights on their functions. Gene expression data from several species show that at least one GID1 gene is expressed in every sampled tissue, with a strong bias of GID1b expression towards underground tissues and dry legume seeds (typically with low GA levels). Taken together, our results support that GID1ac retained canonical GA signaling roles, whereas GID1b specialized in conditions of low GA concentrations. We propose that this functional specialization occurred initially at the gene expression level and was later fine-tuned by specific mutations that conferred greater GA affinity to GID1b, including a Phe residue in the GA-binding pocket. Finally, we discuss the importance of our findings to understand the diversification of GA perception mechanisms in land plants.

## INTRODUCTION

Gibberellins (GAs) are hormones that regulate various processes in plant development, particularly during seed germination, flowering, pollen development and stem elongation (Olszewski et al. 2002). The classic GA signaling pathway is characterized by the recognition of bioactive GA (e.g. GA3 and GA4) by the receptor GIBBERELLIN INSENSITIVE DWARF1 (GID1). GID1 is a nucleocytoplasmic protein (Livne and Weiss 2014) that was initially identified in rice (OsGID1, *Oryza sativa*) (Ueguchi-Tanaka et al. 2005). Upon interaction with GA, GID1 undergoes a conformational change that increases its affinity for DELLA, proteins that typically inhibit GA signaling by: interacting and blocking the activity of transcription factors that drive GA transcriptional programs (Murase et al. 2008); working as co-activators of negative regulators of GA signaling and; recruiting chromatin remodeling proteins to promoter regions (Nelson and Steber 2016). In the canonical GA signaling pathway, the GA-GID1-DELLA complex is recognized by the SCF^SLY1^ ubiquitin ligase, which ubiquitinates DELLA proteins, promoting their proteasomal degradation (Dill et al. 2004; Fu et al. 2004; Gomi et al. 2004; McGinnis et al. 2003; Peng et al. 1997). Therefore, the down-regulation of DELLA is the process that ultimately trigger the classic GA effects (Fleet and Sun 2005). Alternative GA signaling pathways have also been proposed, such as a GA-independent (GID1-mediated) (Yamamoto et al. 2010) and DELLA-independent pathways (Fuentes et al. 2012). Interestingly, both the canonical and alternative pathways described above rely on GID1, which appears to have a central role in GA signaling.

GID1 receptors evolved from a larger family of Hormone Sensitive Lipases (HSLs). Comparison of HSLs with the rice GID1 revealed important differences: the His from the HSL catalytic triad (Ser-Asp-His) is replaced by Val in GID1; the last Gly of the HGGG motif is substituted by Ser in GID1 and; the extensive divergence between N-terminal lid of GID1 and HSLs (Hirano et al. 2012). Detailed structural analyses of the GA-GID1a-DELLA complex support that these changes are critical for GA binding. Other GID1a amino acid residues were also found to be involved in GA interaction: Gly^114^, Gly^115^, Ser^116^, Ile^126^, Tyr^127^, Ser^191^, Phe^238^, Val^239^, Asp^243^, Arg^244^, Tyr^247^, Gly^320^, Tyr^322^, Leu^323^ (core domain residues) and; Ile^24^, Phe^27^, Lys^28^, Tyr^31^, Arg^35^ (N-terminal extension residues) (Murase et al. 2008).

Three GID1 receptor genes have been characterized in *Arabidopsis thaliana* (GID1a, GID1b and GID1c). Although some level of functional redundancy was found between these genes, each of them apparently play specific roles in different developmental stages (Griffiths et al. 2006; Iuchi et al. 2007; Suzuki et al. 2009; Willige et al. 2007). GID1 receptors were also characterized in several other plants, such as ferns (Hirano et al. 2007), cotton (Aleman et al. 2008), barley (Chandler et al. 2008) and wheat (Li et al. 2013). A previous phylogenetic reconstruction of GID1 receptors uncovered the presence of three major groups: eudicot GID1ac, eudicot GID1b and monocot GID1, supporting that a diversification of this family occurred after the split of monocots and dicots (Voegele et al. 2011). In addition to the phylogenetic separation of GID1ac and GID1b subfamilies, a number of important features related to the functional specialization of GID1 subfamilies have been described: 1) a remarkable difference in their transcriptional profiles across several tissues, such as in roots (Griffiths et al. 2006) and during germination (Bellieny-Rabelo et al. 2016); 2) the transcriptional down-regulation of GID1ac, but not GID1b, by GA. (Voegele et al. 2011); 3) The different affinity of GID1 subfamilies for GA, with GID1b displaying greater affinity for GA_3_ and GA_4_ than GID1a and GID1c (Nakajima et al. 2006) and; 4) The preference of specific GID1 proteins for particular DELLA groups (Hirano et al. 2007), potentially increasing the complexity involved in GA signaling.

Although important aspects of the GID1 family have been elucidated since its discovery and structural determination, important questions remain to be answered regarding GID1 expansion and diversification, the distribution of GID1ac and GID1b subfamilies in major eudicot lineages and the major evolutionary forces shaping the eudicot subfamilies at the sequence and expression levels. Here we performed a comprehensive survey of GID1 proteins in 52 plant genomes and integrate this data with protein structure and gene expression data. Our results provide important insights on the evolutionary history of the GID1 family in land plants, including findings such as: 1) a detailed phylogenetic reconstruction of GID1s and the identification of the main expansion and diversification events, including a potential contribution of whole-genome duplication (WGD) events to the structure of the GID1 family in eudicots; 2) the conservation and divergence of key amino acid residues involved in GA and DELLA binding by GID1b and GID1ac subfamilies and; 3) the important contribution of gene expression divergence in the establishment and divergence of the GID1ac and GID1b subfamilies in eudicots. Finally, we discuss theoretical aspects regarding the evolution of GA perception mechanisms, which that can fuel future computational and experimental studies.

## RESULTS AND DISCUSSION

### Expansion and diversification of GID1 receptors in major groups of land plants

A total of 52 diverse plant genomes, including angiosperms, gymnosperms and basal land plants (i.e. a lycophyte and a bryophyte) (Supplementary Table S1), were screened for GID1 proteins (see methods for details). Due to their high sequence similarity to HSLs, GID1s were separated with the aid of a phylogenetic reconstruction strategy (see methods for details) (Supplementary Fig. S1). We identified a total of 138 GID1 genes, with a median of two GID1s per genome (Supplementary Table S2) and ~81% of the angiosperms containing 2-3 GID1 genes (Fig. 1). All eudicots except the early-branching *Aquilegia coerulea* have more than one GID1, which were classified using BLAST searches against *Ar. thaliana* GID1s and phylogenetic reconstructions by Bayesian and Maximum Likelihood approaches (see methods for details) (Fig. 2; Supplementary Table S2; Supplementary Fig. S2). The species with the greatest number of GID1s are *Gossypium hirsutum* (nine), *Go. raimondii* (six) and *Glycine soja* (six) in eudicots, and *Musa acuminata* (six) in monocots (Fig. 1). The 138 GID1s can be divided in four statistically supported groups (Voegele et al. 2011): group I (GID1ac) and II (GID1b), both with eudicot sequences; group III, with monocot GID1s and; group IV, containing GID1s from gymnosperms and basal plants. While GID1s from basal land plants and gymnosperms formed a separate small group, angiosperm GID1s diversified in the three former groups (Fig. 2). Our results support the monophyly of all plant GID1s, most likely from a single ancestral gene. The only GID1 from *Amborella trichopoda* (a basal angiosperm) is a sister group of the monocot and eudicot clades (Fig. 2), supporting the expansion and divergence of GID1s after the emergence of angiosperms. More precisely, this diversification process happened after the split of Ranuncales, since *Aq. coerulea* has only one GID1 that is an early branch in the GID1ac clade (Fig. 2). Our results also indicate that the GID1ac group is more closely related to the ancestral GID1, whereas the GID1b subfamily originated in eudicots after the separation of monocots, possibly via the *gamma* polyploidy, a whole genome triplication event shared by all core eudicots (Jiao et al. 2012). Conversely, monocot GID1s form a single clade without apparent strong diversification other than some recent lineage-specific expansions, particularly in banana (discussed below).

**Fig. 1.**
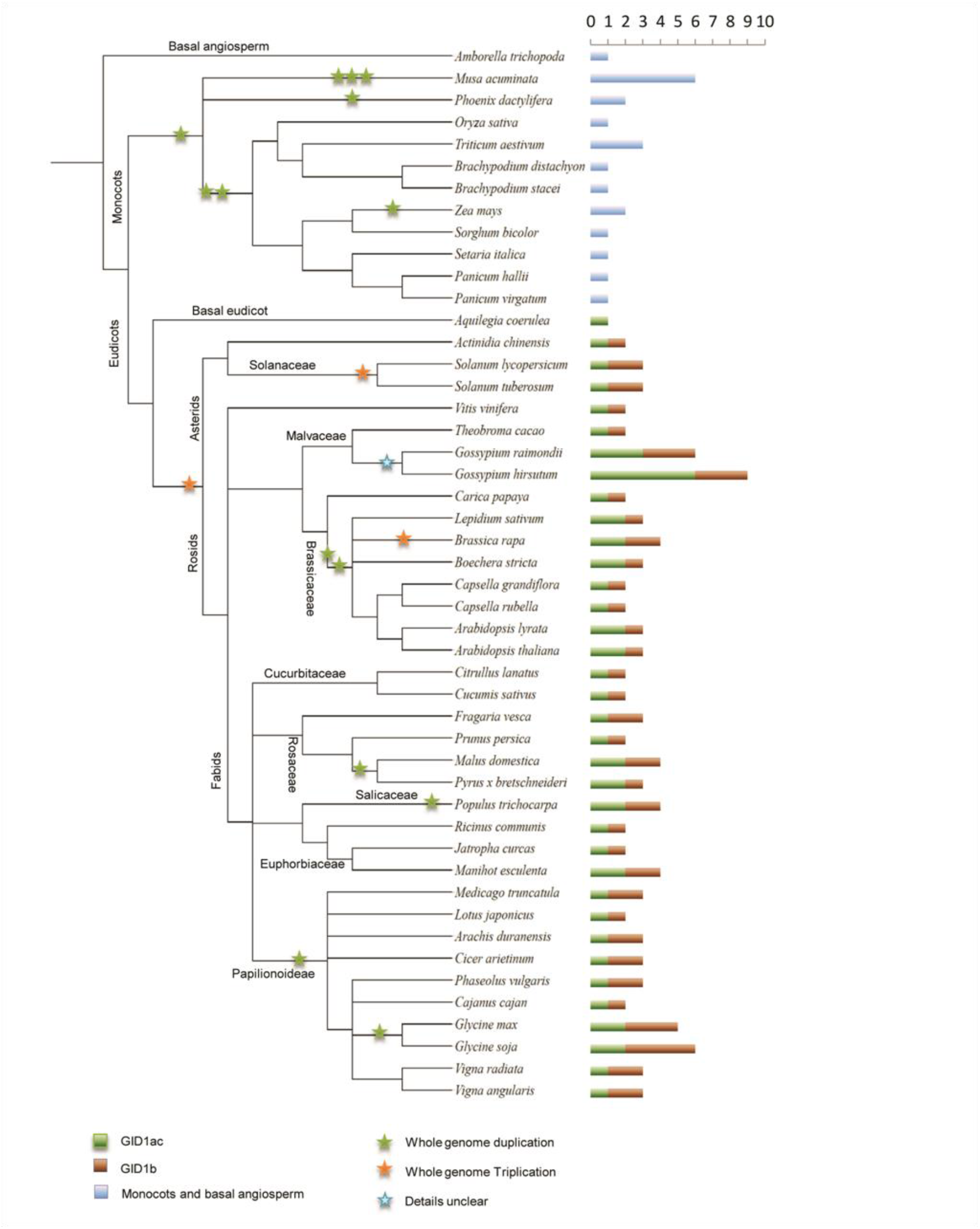
Number of GID1 genes in different angiosperms. GID1 counts in each species are represented as horizontal bars, colored according to subfamily. Polyploidization events are marked with colored stars. Species tree was generated using PhyloT (http://phylot.biobyte.de/). Branch lengths do not represent evolutionary time.

**Fig. 2.**
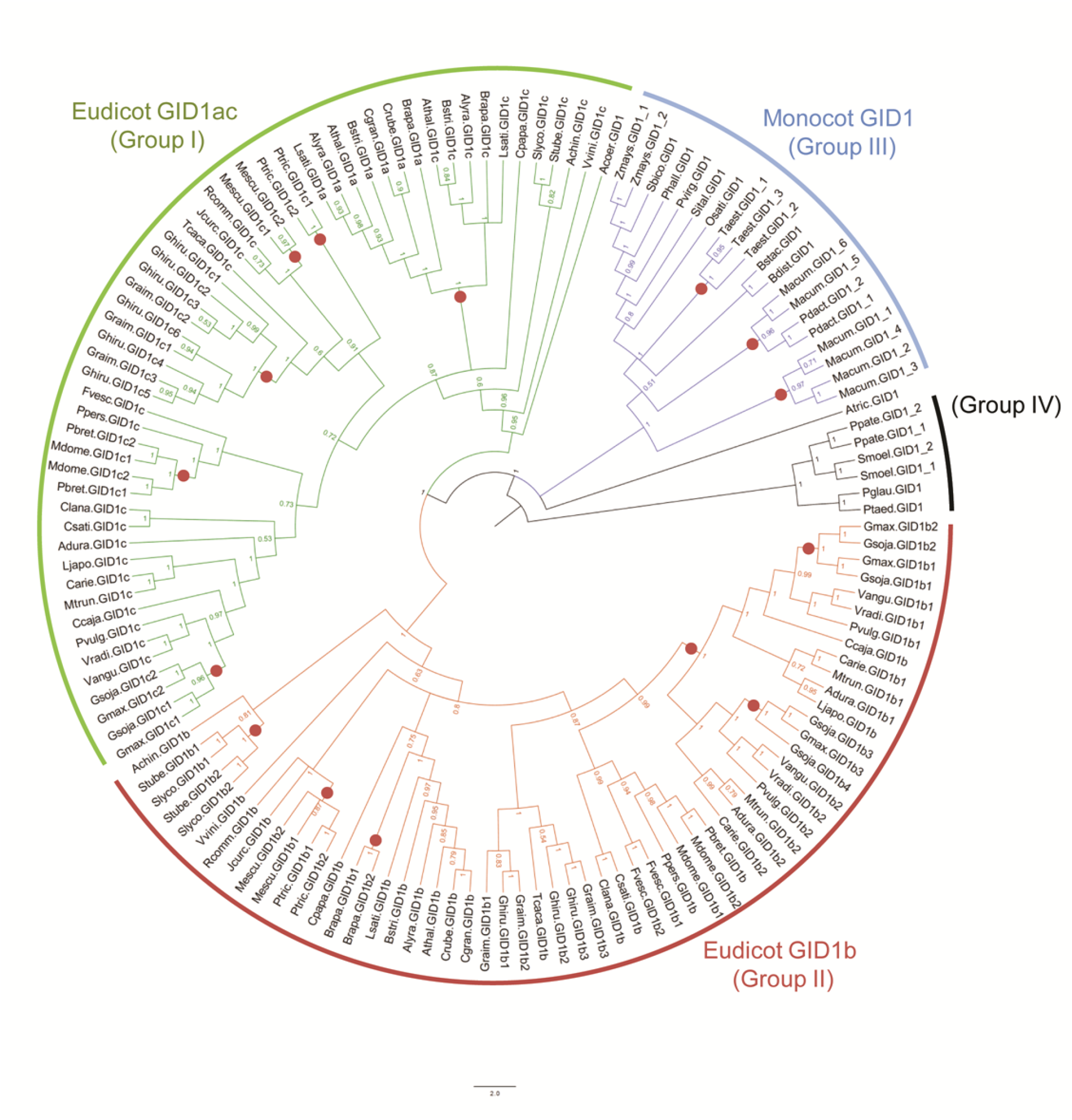
Phylogenetic reconstruction of the 138 GID1 proteins identified in 52 plant species. Multiple sequence alignment of GID1 proteins was carried with PROMALS3D and phylogenetic reconstruction performed with MrBayes. The GID1 proteins were classified in four groups, which are represented in different colors. Red circles show genes potentially originated by whole-genome duplication events.

Next, we sought to explore the evolutionary history of the eudicot GID1ac and GID1b subfamilies. It is clear from our results that there is at least one GID1ac and one GID1b in every core eudicot, implying that these subfamilies acquired important non-redundant roles early in the evolution of eudicots (Voegele et al. 2011). Although GID1ac and GID1b subfamilies have comparable total sizes, their distribution is not uniform across lineages (Fig. 1). Asterids and many rosids have a single GID1ac, although some independent lineage-specific expansions happened after the separation of these two large groups (Fig. 1, Fig. 2). In Malvaceae, the two *Gossypium* species experienced a more recent GID1ac expansion, after the split from *Theobroma cacao*. In the order Brassicales, the GID1ac subfamily expanded after the separation of Brassicaceae and Caricaceae, with the emergence of a well-defined clade (harboring proteins related to *Ar. thaliana* GID1a) in the former, whereas *Carica papaya* preserved a single GID1c, outside of the GID1a clade (Fig. 2). In fact, all GID1a proteins belong to a Brassicaceae-specific monophyletic clade nested inside GID1c; this GID1a clade could have emerged at the WGD events that took place after the split of Brassicaceae and Caricaceae (Schranz 2006). Interestingly, with the exception of *Capsella grandiflora* and *Capsella rubella* (Fig. 2), the Brassicaceae species retained both GID1a and GID1c genes, indicating that they also play non-redundant roles (Suzuki et al. 2009). Nevertheless, it has been shown that GID1a and GID1c can compensate the absence of each other during *Ar. thaliana* seed germination (Voegele et al. 2011), suggesting that such non-overlapping roles are performed in other conditions/tissues (Griffiths et al. 2006). *Capsella species* are the only core eudicots without a classical GID1c, indicating a displacement of GID1c by GID1a in this genus. Therefore, these species would be good models to study the recent functional diversification within the GID1ac clade. Other GID1ac duplications that could be attributed to WGD events were also found in Salicaceae (*Populus trichocarpa*), *Glycine*, *Manihot esculenta* and in the most recent ancestor of *Malus domestica* and *Pyrus x bretschneideri* (Fig. 2). Further, all the Fabaceae species except *Gl. max* and *Gl. soja* have a single GID1ac. Our phylogenetic analysis indicates that one of the GID1ac paralogs was rapidly lost after the legume WGD and the remaining GID1ac gene was later duplicated at the *Glycine* WGD. This scenario is also supported by the presence of gene pairs with low Ks values the domesticated and wild soybeans (Fig. 2, Supplementary Table S3).

On the other hand, GID1b is mainly expanded in legumes, most likely due to the WGD events that happened at the base of Papilionoideae and *Glycine*. Except for *Lotus japonicus* and *Cajanus cajan* (which independently lost one GID1b paralog), all other legumes retained duplicated GID1b sets, with two duplication rounds accounting for the 3-4 GID1b genes found in soybeans (Fig. 1, Fig. 2). Similarly to what was observed for GID1ac, there is also a soybean GID1b pair (*Gmax.GID1b1* and *Gmax.GID1b2*) that probably originated in the Glycine WGD (Fig. 2). This gene pair has a low Ks value (i.e. 0.187) that is compatible with the Glycine WGD age. Although these genes are not located in previously identified large homeologous segments (Severin et al. 2011), they show some level of conservation in their genomic neighborhood (Supplementary Fig. S3). Further, this scenario implies a loss of one GID1b in *Gl. max* after the separation from *Gl. soja*, possibly during soybean domestication; this hypothesis is supported by phylogenetic reconstructions (Fig. 1) and the low Ks values of the respective surviving *Gl. soja* paralogous pair (*Gsoja.GID1b3* and *Gsoja.GID1b4*; Ks = 0.127; Fig. 2, Supplementary Table S3). Other expansions of GID1b genes can also be found in *Manihot esculenta, Fragaria vesca, Populus*, *Solanum* and *Gossypium*, for which polyploidization events have been documented or predicted (Mühlhausen and Kollmar 2013; Sato et al. 2012; Tuskan et al. 2006; Xu et al. 2011; Zhang et al. 2015). Conversely to what was observed in GID1ac, the GID1b subfamily size is constrained in Brassicales, in which only *Brassica rapa* has more than one gene, which may have originated by a recent *Brassica* whole-genome triplication event (Fig. 1, Fig. 2). Remarkably, even after two WGDs, Brassicaceae species have a single GID1b, indicating that the retention of GID1b duplicates is peculiar to a few clades, particularly legumes.

In monocots we have not found large and diversified GID1 subgroups (Fig. 1, Fig. 2), although there are multiple recent duplications in various lineages (e.g. maize and wheat). The most striking expansion of GID1 in monocots occurred in banana, in which six GID1s were found (Fig. 1, Fig. 2). Interestingly, although three recent WGDs have been identified in the banana genome (D’Hont et al. 2012), the Ks values of these GID1 pairs are far greater than expected for duplicates generated in these WGDs (Supplementary Table S3). The only banana GID1 pair with low Ks, *Macum.GID1_2* and *Macum.GID1_3*, is separated by less than 20 kb, with a single intervening gene, supporting an origin via proximal (i.e. tandem) duplication. Furthermore, these banana GID1 pairs are outside of the homeologous blocks identified in the banana genome project (D’Hont et al. 2012), suggesting that they originated by duplication mechanisms other than the recent WGD.

### GID1 intron-exon structure is largely conserved throughout the evolution of land plants

In addition to genomic locations and phylogenetic reconstructions, we also investigated the GID1 gene architectures (i.e. intron-exon structures) and intron phases (Fig. 3, Supplementary Fig. S4). There are three possible intron phases: phase 0, in which an intron is located between two codons; phase 1 and 2, with introns between the first and second, and between the second and third codon nucleotides, respectively. This analysis revealed a strong conservation at the level of intron-exon structure. We found that 106 out of 126 angiosperm GID1s (~84 %) with available gene structure have the same basic gene structure, comprising a short and a long exon (average length of 42 bp and 990 bp, respectively), with an intervening phase 0 intron of ~610 bp (Fig. 3; Supplementary Fig. S4). Gene structure conservation is even greater in eudicots, which have 95 out of 108 genes (88%) with the canonical architecture (Fig. 3). Remarkably, gene structure conservation in eudicots is independent of subfamily division, strongly supporting the evolution of eudicot GID1 subgroups from a single ancestor, most likely with the gene structure similar to that of *Acoer.GID1*. The canonical GID1 gene structure is also largely preserved in monocots, although three different architectures are found in banana GID1s (Fig. 3). Importantly, the lycophyte GID1s resemble this architecture, indicating that it represents an ancestral state that has been widely conserved throughout angiosperms. However, gymnosperm and bryophyte GID1 gene architectures are distinct from this theme, as are those of other few angiosperm GID1s.

**Fig. 3.**
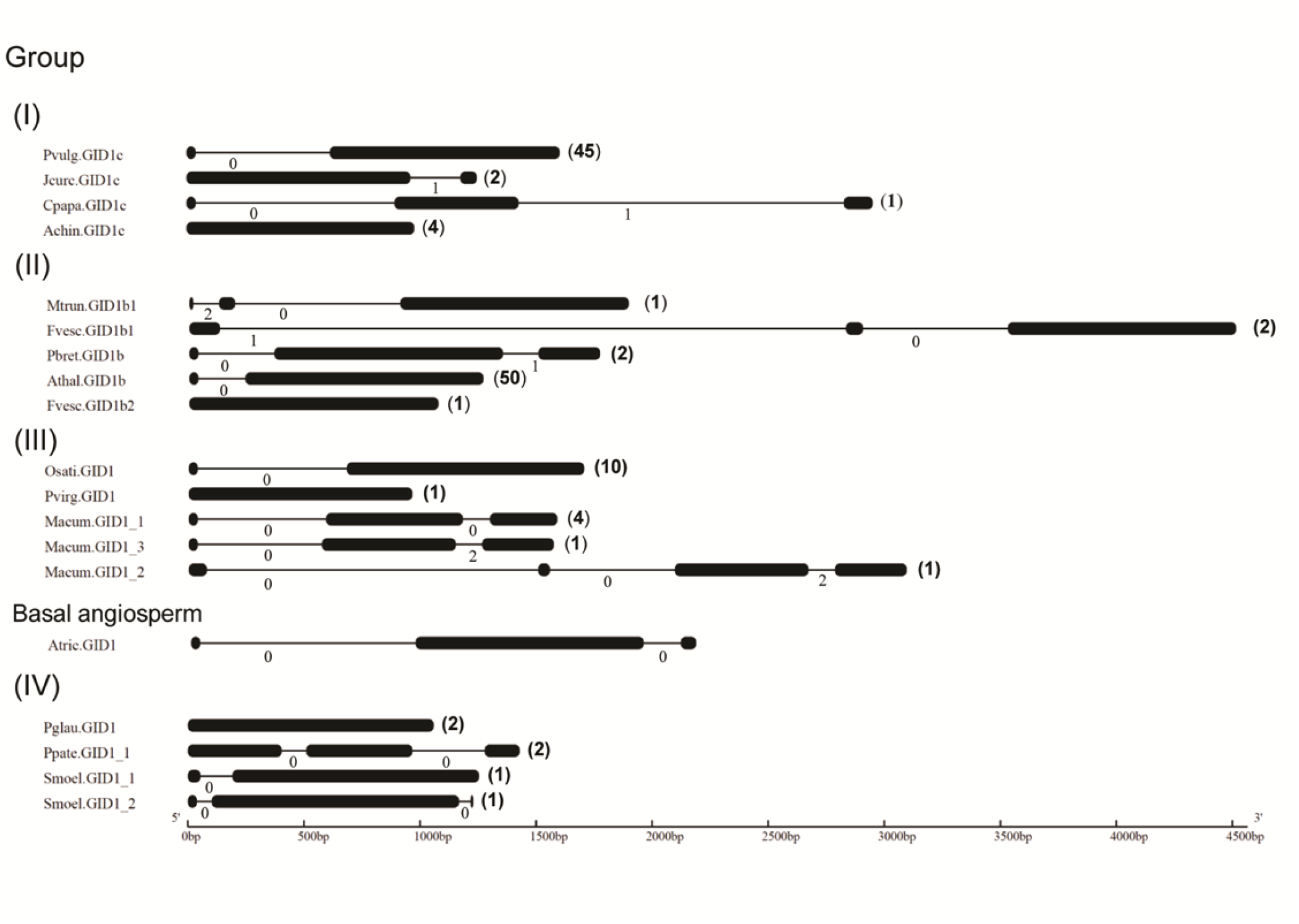
Diagram of representative GID1 intron-exon architectures. Thin lines and thick bars represent introns and exons, respectively. Numbers below introns and at the right side of the gene architectures represent intron phases and number of occurrences of each structure, respectively. For comparison purposes, the intron-exon structure of the *Am. trichopoda* GID1, a basal angiosperm, is shown below the gene structures of monocot GID1s.

### GID1 subfamilies have shared and specific structural features

Two critical steps in the evolution of GID1s from the HSL family were the loss of catalytic activity and the emergence of GA-binding properties (Hirano et al. 2012). We performed extensive sequence comparisons to better understand the conservation and divergence of GID1 subfamilies. Notably, the characteristic motifs HGGS and GDSSG are conserved in all analyzed GID1s, except for the presence of HGGG instead of HGGS and GDSAG instead of GDSSG in bryophytes) (Fig. 4). Moreover, a SUMO-Interaction Motif (SIM; amino acids W[V/I]LI), that is important for the recognition of SUMOylated DELLA proteins (Conti et al. 2014; Nelis et al. 2015), is also conserved across GID1s, again except in bryophytes. Five GID1s have amino acid substitutions in the first position of the SIM: Met (in Bdist.GID1 and Bstac.GID1), Tyr (in Mtrun.GID1b) and Phe (in Pgalu.GID1 and Ptaed.GID1). A hydrophobic surface in the N-terminal lid (Leu^18^, Trp^21^, Val^29^, Ile^33^, Leu^45^ and Tyr^48^ in *Ar. thaliana* GID1a) forms a DELLA-binding surface (Murase et al. 2008; Shimada et al. 2008) and is also highly conserved in almost all GID1s (Fig. 4, Supplementary Fig. S5); we mapped these hydrophobic residues in the alignment and found that Leu^45^ is fully conserved, whereas the remaining positions tolerate substitutions by other hydrophobic residues (Fig. 4, Supplementary Fig. S5). For example, instead of Ile^33^, Met^33^ is present in all monocots (except rice) and in very few eudicots, mainly Solanaceae (i.e. Slyco.GID1b1, Slyco.GID1.b2, Stube.GID1.b1, Stube.GID1b2 and Achin.GID1b). Met^33^ seems to be the ancestral state, as it is present in *Am. trichopoda* and *Aq. coerulea*. Further, because these species have only a single GID1, we hypothesize that Met^33^ can be part of GID1 DELLA binding surfaces. Unexpectedly, some GA interacting residues (i.e. Asn^218^, Phe^238^, Val^239^, Asp^243^, Arg^244^, Tyr^247^ in *Ar. thaliana* GID1a) are missing in all banana GID1s (Supplementary Fig. S5C). The impact of these mutations in the banana GID1s warrants further investigation, for example by expressing banana GID1s in rice GID1 mutants.

**Fig. 4.**
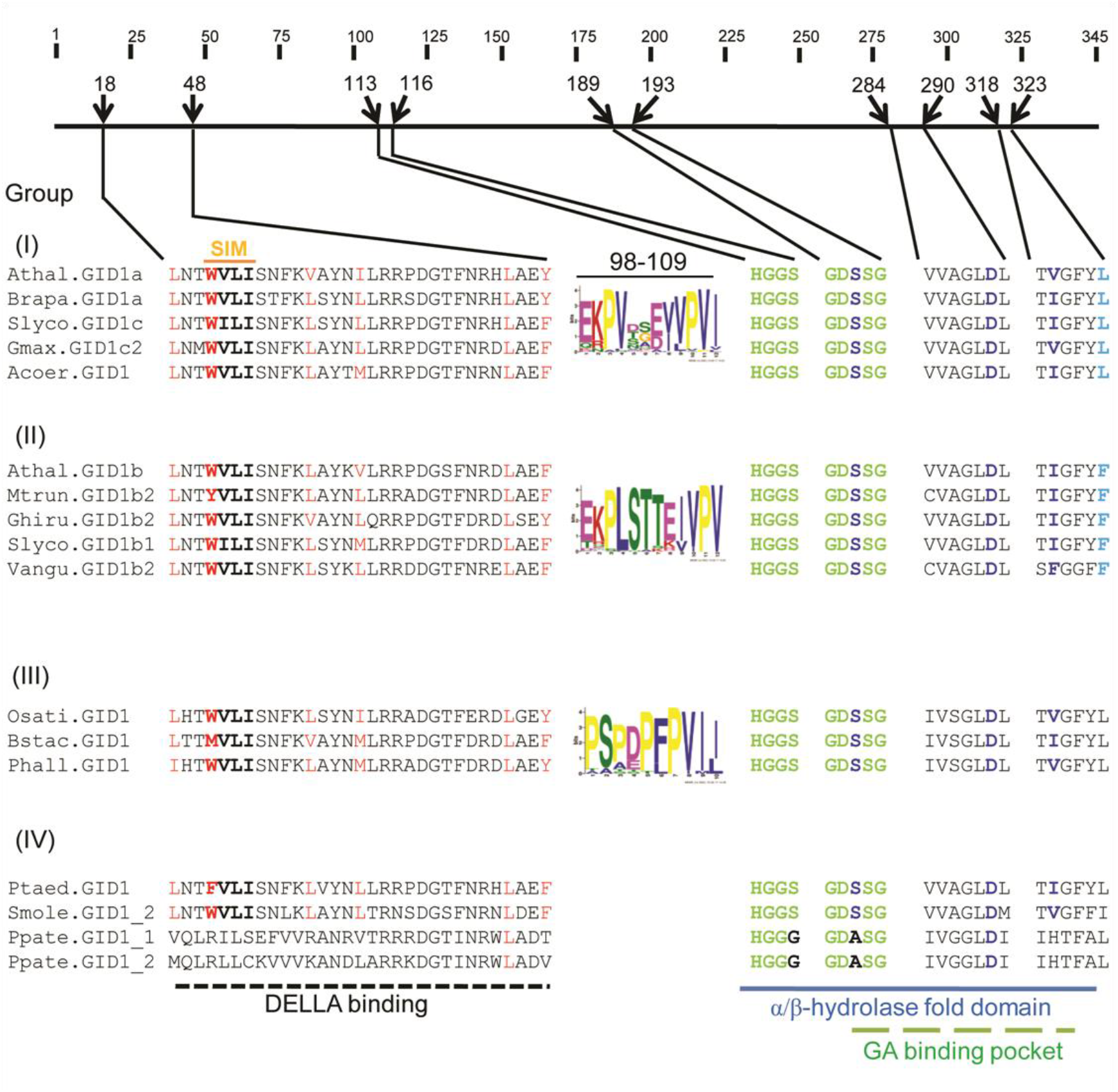
Schematic representation of archetypal GID1s. The thick line with coordinates represents the positions of important GID1 features, using the *Ar. thaliana* GID1a as a reference. HGGS and GDSSG motifs are shown in green; the SIM motif (W[V/I]LI) is also tagged. DELLA binding residues are shown in red and the ‘catalytic triad’ involved in binding GA (Ser, Asp, and Val/Ile) are highlighted in dark blue. Subgroup specific motifs are represented with logos. One functionally divergent site between GID1ac and GID1b, which is also a GA interacting residue, is marked with sky blue color. All functionally divergent sites can be seen in Supplementary Fig. S5 and Table 1.

Although GID1s display an overall high level of sequence similarity, we were able to clearly define four major clades (Fig. 1), which support some level of functional divergence between them. To better understand the conservation patterns in the family, we sought to analyze conserved and specific motifs in GID1 subfamilies (Supplementary Fig. S5). We found five motifs that are conserved in all four clades (except in bryophyte GID1s, group IV). Three of those were well known motifs: Motif 1, which encompasses the SIM, GA- and DELLA-interacting residues; Motif 3, which contain the HGGS motif and; Motif 4 harbors the GDSSG domain and GA interacting residues. The remaining two motifs are Motif 5 and 6, which harbor other GA-binding residues (Supplementary Fig. S5). We also identified motifs specific to the GID1ac (Motif 2), GID1b (Motif 7) and monocot GID1s (Motif 8) sub-groups (Fig. 4, Supplementary Fig. S5). Further, these three motifs correspond to the same alignment region and their within-group conservation patterns suggest that they might play important subfamily-specific roles.

To further explore the mechanistic differences of GID1ac and GID1b, we have also predicted functionally divergent sites using three different programs (see methods for details). A total of nine alignment positions were predicted to be functionally divergent between GID1ac and GID1b groups (Table 1). We mapped these residues on the tertiary structure of the Athal.GID1a (Fig. 5), as well as on the predicted conserved motifs described above. Two sites, Asp^102^ and Gly^103^ in Athal.GID1a (Ser^102^ and Thr^103^ in Athal.GID1b) were inside the specific motifs discussed above (Fig. 4, Supplementary Fig. S5). Interestingly, the positions 102 and 103 are much more conserved in the GID1b (Ser^102^ and Thr^103^ in Athal.GID1b) than in the GID1ac subfamily (Fig. 4, Supplementary Fig. S5), supporting that these sites are under type I functional divergence (Gu 2001, 1999) (Table 1). Four other functionally divergent sites were highly conserved within GID1ac and GID1b subgroups but with important amino acid changes (e.g. Leu^323^ in GID1ac subfamily and Phe^323^ in GID1b subfamily) between them, suggesting type II functional divergence (Table 1) (Gu 2001, 1999).

**Fig. 5.**
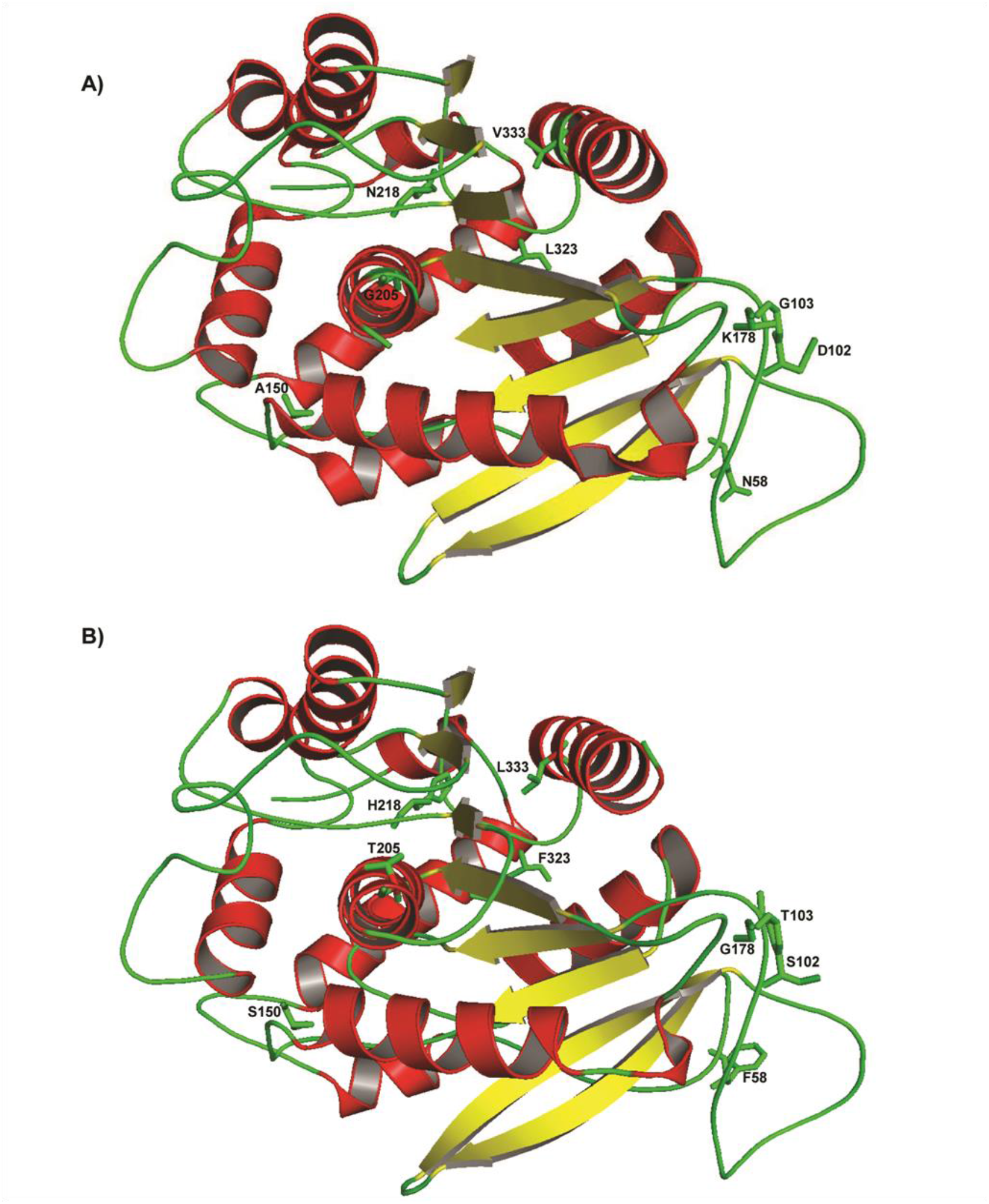
Localization of critical amino acids in the 3D structures. Functionally divergent sites were mapped on the crystal structure (as sticks) of the native (A) and mutated (*in silico*) (B) *Ar. thaliana* GID1a (2ZSH).

**Table 1.** Number of sites predicted to be functionally divergent in GID1ac and GID1b groups.

| Position in GID1a<br>crystal structure | GID1ac group (Athal.GID1a<br>as reference) | GID1b group (Athal.GID1b<br>as reference) | Functional Divergence<br>Type |
| --- | --- | --- | --- |
| **Loop** | Asn<sup>58</sup> | Phe<sup>58</sup> | Type I |
| **Loop** | Asp<sup>102</sup> | Ser<sup>102</sup> | Type I |
| **Loop** | Gly<sup>103</sup> | Thr<sup>103</sup> | Type I |
| **Loop** | Ala<sup>150</sup> | Ser<sup>150</sup> | Type II |
| **Loop** | Lys<sup>178</sup> | Gly<sup>178</sup> | Type I |
| **α3 helix** | Gly<sup>205</sup> | Thr<sup>205</sup> | Type U |
| **Loop** | Asn<sup>218</sup> | His<sup>218</sup> | Type II |
| **η2 helix** | Leu<sup>323</sup> | Phe<sup>323</sup> | Type II |
| **α7 helix** | Val<sup>333</sup> | Leu<sup>333</sup> | Type II |

Intriguingly, one of the functionally divergent sites, Leu^323^ (in GID1ac; Phe^323^ in GID1b and Leu^330^ in rice GID1), is involved in hydrophobic interactions with GA (Murase et al. 2008). Previous studies in rice demonstrated that mutation of GA interacting residues, including the substitution of Leu^330^ for Ile^330^ or Ala^330^, reduced the GID1 affinity and specificity for GAs (Hirano et al. 2007; Shimada et al. 2008; Xiang et al. 2011). We performed *in silico* mutagenesis with FoldX to estimate the effects of converting Leu^323^ into Phe^323^ on the GA binding pocket of Athal.GID1a (2ZSH and 2ZSI), followed by docking analysis of mutated 2ZSH with GA_3_ and mutated 2ZSI with GA_4_. The native and mutant docked GID1-GAs had similar hydrogen bond lengths (Murase et al. 2008). In previously reported structures, the O7-2 atom of GA_3_/GA_4_ formed a hydrogen bond to the Oγ atom of Ser^191^ with a distance of 2.9 Å (for GA_3_) and 3.2 Å (for GA_4_) (Fig. 6A, Fig. 6C). These corresponding distances became longer (3.5 Å for both GA_3_ and GA_4_), although within the range of hydrogen bonds (Fig. 6B, Fig. 6D). Interestingly, we found that Phe is closer to GA_3_/GA_4_ than Leu with a significant difference of ~1Å, suggesting that Phe^323^ in GID1b confers a tighter binding pocket that could be related with the higher affinity of GID1b for GA_3_/GA_4_. Interestingly, this higher affinity of GID1b has been attributed to a partially closed configuration of the N-terminal lid. We hypothesize that Phe^323^ may also contribute to this phenomenon.

**Fig. 6.**
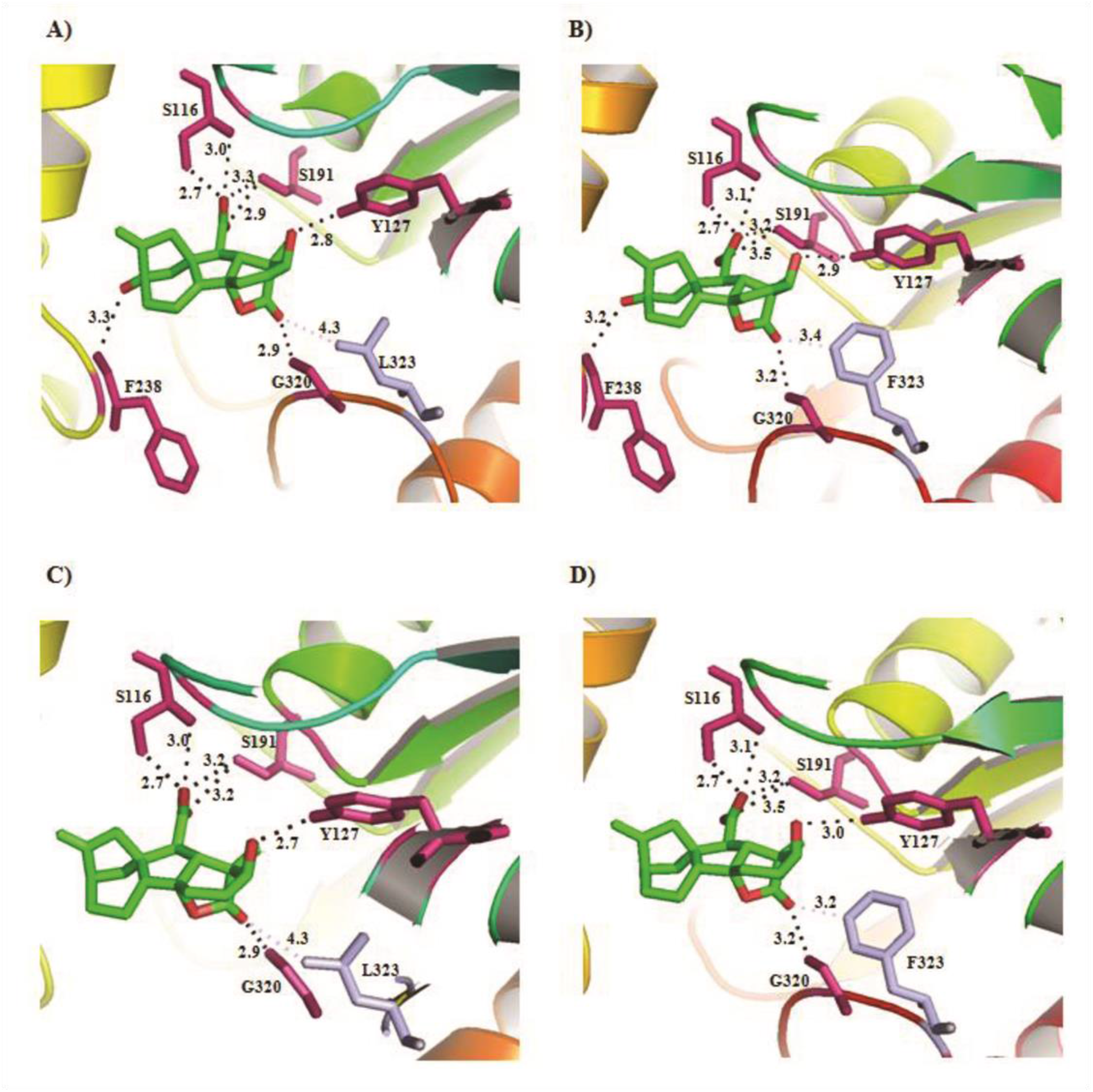
Comparison of GID1a-GA in the native versus mutated GID1a-GA. **A**) native structure of 2ZSH-GA_3_; **B**) mutated structure of 2ZSH-GA_3_; **C**) native structure of 2ZSI-GA_4_ and; **D**) mutated structure of 2ZSH-GA_4_. Amino acid Leu^323^ of GID1a and its corresponding amino acid in GID1b (Phe^323^) are shown in light blue; distances of these amino acids to GA_3_/GA_4_ are also shown in dotted light blue. Hydrogen bonds are shown in dotted black lines.

### GID1 subfamilies have substantial divergence in their expression patterns

Given the expansion and diversification of GID1 subfamilies, we sought to study their expression profiles as a means to understand their functional specialization. In *Phaseolus vulgaris*, we found a general higher GID1 expression, particularly of GID1b, in underground tissues (i.e. roots and nodules). Further, the expression variance between tissues is greater in GID1b (*Pvul.GID1b1* and *Pvul.GID1b2*) than in GID1c (*Pvul.GID1c*), probably due to the absence of GID1c paralogs (Supplementary Fig. S6). In soybean, we also found a remarkable activation of a GID1b paralog (*Gmax.GID1b3*) in roots and nodules, in addition to a conspicuous expression peak in in flowers (not observed in common bean) (Supplementary Fig. S7). Interestingly, the homeolog of *Gmax.GID1b3* that originated in the *Glycine* WGD was lost in *Gl. max* (but not in *Gl. soja*) (Fig. 2), possibly during domestication. We speculate that the specialized expression profile of *Gmax.GID1b3* and the lack of a close homeolog may be involved in the selection of traits of agricultural interest. Interestingly, our group has shown that soybean GID1b genes are highly expressed in the embryonic axes of dry seeds, being down-regulated as germination proceeds, a trend that is opposite to that of GID1c genes (Bellieny-Rabelo et al. 2016) (Supplementary Fig. S8). This scenario can be part of a system to detect low GA levels and trigger important signaling processes until the canonical GID1c-mediated GA signaling pathway is activated in the onset of germination (Bellieny-Rabelo et al. 2016; Griffiths et al. 2006; Hauvermale et al. 2015; Nakajima et al. 2006). We have also analyzed the expression of GID1 genes in a third legume species, *Me. truncatula* (Supplementary Fig. S9). Similarly to what was observed in soybean and common bean, GID1b is also more expressed than GID1c in most *Me. truncatula* tissues and at least one GID1b gene is highly expressed in roots and nodules. Interestingly, *Mtrun.GID1b1* transcripts accumulate during seed maturation, whereas *Mtrun.GID1c1* transcription is reduced, a trend that is similar to what was observed in soybean (Bellieny-Rabelo et al. 2016), but not in *Ar. thaliana* (Voegele et al. 2011). Taken together, these results indicate that the expression divergence of GID1b and GID1ac in seed development and germination occurred before the split of soybean and *Medicago* [~52 MYA (Kumar et al. 2017)], but after the origin of Rosids. Nevertheless, this scenario will be clearer only when more gene expression data of GID1 genes during seed development and germination become available for other species.

Similarly to legumes, *Athal.GID1b* is more expressed than *Athal.GID1a* and *Athal.GID1c* in roots, whereas GID1ac expression is dominant in leaves, flowers and developing seeds (Supplementary Fig. S10). Thus, our results indicate that the specialization of GID1b towards roots, nodules and dry seeds support the scenario where GID1b, probably because of its higher affinity for GA, is particularly suitable for conditions of low GA concentrations and/or tissues with high GA sensitivity (Tanimoto 1987, 1994). It has been shown that GA regulates root elongation and thickening (Tanimoto and Hirano 2013). In root elongation, GA action specifically takes place at the endodermis (Ubeda-Tomás et al. 2008). In addition, GA also influences the number and length of root meristems (Tanimoto and Hirano 2013; Ubeda-Tomás et al. 2009). Thus, GID1b probably specialized to mediate GA signaling in eudicot roots in the presence of low hormone concentrations

We have also investigated GID1 expression in monocots and in the lycophyte *Se. moellendorffii*, in which there are often fewer GID1 genes and no family subdivision (Supplementary Fig. S11, S12 and S13). Interestingly, we found that GID1 is highly expressed in all tissues, with at least one GID1 gene expressed in high levels in roots. Collectively, our results show that the high expression of GID1 in roots dates back to the origin of the canonical GA perception system in lycophytes (Hirano et al. 2007; Nelson and Steber 2016), far earlier than the emergence of seed plants. In species without GID1 subfamilies (e.g. monocots and lycophytes), often harboring a single GID1 gene, this gene is expressed in almost all tissues and displays high expression in roots. We hypothesize that after the divergence of GID1ac and GID1b subfamilies, the former retained roles more related to the ancestral GA perception system (already present in lycophytes), and was later recruited to more modern features like seed germination. On the other hand, GID1b specialized in conditions of low GA concentrations (e.g. roots and germinating legume seeds) through biased gene expression and accumulation of mutations that increased its affinity for GA (Nakajima et al. 2006). Further, with GID1ac mediating canonical GA signaling, GID1b was also free to integrate alternative GA perception mechanisms, such as GA-independent DELLA binding and non-proteolytic GA signaling (Fuentes et al. 2012; Yamamoto et al. 2010).

Important aspects regarding the origin of GA perception system remain to be elucidated. While the lycophyte *Se. moellendorffii* and the bryophyte *Physcomitrella patens* have the key components of the canonical GA perception machinery, such system seems to be absent in the latter (Hirano et al. 2007; Nelson and Steber 2016), which is supported by several lines of evidence (Hayashi et al. 2010; Vesty et al. 2016; Yasumura et al. 2007), such as: 1) *Physcomitrella patens* GID1 and DELLA do not interact; 2) *Physcomitrella patens* GID1 does not interact with GA; 3) DELLA-deficient *P. patens* strains do not exhibit derepressed growth like that observed in DELLA-deficient angiosperms; 4) *Physcomitrella patens* DELLA does not suppress GA response in rice, although it can repress growth in *Ar. thaliana*. On the other hand, certain bioactive diterpene hormones from early steps of the GA biosynthesis pathway (e.g. *ent*-kaurene) promote spore germination in *Physcomitrella patens* (Hayashi et al. 2010; Vesty et al. 2016). Interestingly, ABA can inhibit *Physcomitrella patens* spore germination, strongly supporting the existence of a diterpene/ABA signaling module before the emergence of vascular plants, although apparently not as prominent as that found in seed plants (Hayashi et al. 2010). The key genes involved in diterpene perception in *Physcomitrella patens* remain to be elucidated and could involve direct diterpene recognition by GRAS domain proteins (e.g. DELLA), which were already diversified early in the evolution of land plants (Zhang et al. 2012).

## MATERIALS AND METHODS

### Identification of GID1 proteins in angiosperms

To identify the GID1 proteins in land plants, predicted proteins of 47 angiosperms, two gymnosperms, one lycophyte and one bryophyte were downloaded from various sources (Supplementary Table 1). GID1 homologs were identified in four steps: 1) BLASTP (Altschul et al. 1997) searches using experimentally characterized GID1s from *Ar. thaliana, Lepidium sativum* and rice to search the predicted proteomes of each species (a total 1,995,759 proteins), with e-value and similarity thresholds of ≤ 1e^−5^ and ≥ 38%, respectively. This step resulted in a total of 252 proteins; 2) Only the 238 sequences with the conserved motifs HGG and GXSXG, also shared with HSLs and other plant carboxylesterases (Ueguchi-Tanaka et al. 2005; Voegele et al. 2011), were retained; 3) Bona-fide GID1s were separated from plant carboxylesterases using a phylogenomic approach, as follows: carboxylesterases of *Ar. thaliana* (AT5G23530) and rice (ABA92266) (Hirano et al. 2007) were aligned with the 238 GID1 candidates using PROMALS3D (Pei et al. 2008). The phylogenetic reconstruction was performed with FastTree (Price et al. 2010). A total of 138 GID1s clearly clustered in a monophyletic clade (Supplementary Fig. S1) and were separated from carboxylesterases; 4) redundancy was removed with the aid of BLASTCLUST (95% coverage and 95% identity thresholds) (Altschul et al. 1997). These steps allowed us to identify 130 GID1s. We found only one GID1 in *Vitis vinifera*, but two *V. vinifera* GID1 genes AM479851 and AM468374 were available in Genbank and were used. Our collection was supplemented with *Triticum aestivum* and *L. sativum* GID1s (three from each) (Li et al. 2013; Voegele et al. 2011). One GID1 from *Ca. cajan* was excluded because of the absence of a start codon. The coding sequences of the identified GID1s were also searched in their respective genomes using BLASTN with an e-value threshold of ≤ 1e^−6^ (Altschul et al. 1997), which allowed us to identify an additional *Gl. soja* GID1.

We have not found GID1 genes in the *Picea glauca*, probably due to genome incompleteness. One of identified GID1 genes of *Se. moellendorffi* was fragmented. We obtained GID1 sequences from *Pi. glauca* and *Se. moellendorffi* from individual Genbank entries [Pi. *glauca* (Genbank: BN001188.1) and *Se. moellendorffii* (Refseq: XP_002993392.1, XP_002993392.1)]. Overall, a total of 138 GID1s were used in the analyses (Supplementary Table S2). Species names were abbreviated by the first letter of genus followed by the four letters of the species name (e.g. Athal corresponds to *Ar. thaliana*) (Supplementary Table S2). Eudicot GID1s were classified in GID1a, GID1b and GID1c using *Ar. thaliana* GID1s as reference. Non-eudicot GID1s were simply numbered, as there is no subfamily division in these species.

### Sequence analysis and phylogenetic reconstruction

Multiple sequence alignment of GID1 proteins was carried out using PROMALS3D (Pei et al. 2008) and visually inspected with Jalview (Waterhouse et al. 2009). Large N- and C-terminal gaps were removed. Conserved motifs were analyzed with MEME (Bailey et al. 2009). Phylogenetic reconstructions were performed with MrBayes (v3.2.2; 3,000,000 generations, sampling frequency: 100) (Ronquist and Huelsenbeck 2003). Convergence was assessed using Tracer (v1.6) (http://beast.bio.ed.ac.uk/Tracer) and posterior probabilities were estimated by removing the burn-in generations as required. Alternatively, we have also reconstructed phylogenies using RAxML (best fit model, 100 bootstrap samples) (Stamatakis 2014). Gene structure analysis was performed using GSDS (v2) (Hu et al. 2015). The aligned proteins were used to guide the conversion of cDNA into codon alignments by PAL2NAL (Suyama et al. 2006). CODEML (PAML version 4.9b) (Yang 1997) was used in for Ks calculation. The Goldman and Yang ML method and the F3x4 model were applied.

### Functional divergence, *in silico* mutagenesis and docking

We used three different programs to infer functionally divergent sites of GID1ac and GID1b subfamilies (with default options): FunDi (Gaston et al. 2011), GroupSim (Capra and Singh 2008) and Sequence Harmony (Feenstra et al. 2007). We used a threshold of 0.5 (Gaston et al. 2011) to filter the sites identified by FunDi and GroupSim, and the default threshold in Sequence harmony. Only those sites identified by all three programs were considered. An *in silico* mutagenesis approach was performed with FoldX (Schymkowitz et al. 2005), using the wild type crystal structures 2ZSH (GA_3_–GID1a–DELLA) and 2ZSI (GA_4_–GID1a–DELLA), excluding GA_3_, GA_4_ and DELLA. Crystal structures 2ZSH and 2ZSI were downloaded from the PDB database (Berman et al. 2000). Protein-ligand docking was performed using SwissDock (Grosdidier et al. 2011) and ligands (GA_3_ and GA_4_) were selected from the ZINC database (Irwin and Shoichet 2006). All structures were visualized by PyMOL (http://www.pymol.org/).

### Gene expression data

Gene expression data of GID1 genes were obtained from publicly available sources, as following. Soybean: Soybase (http://soybase.org/soyseq/) (Severin et al. 2010) and from a recent manuscript from our group (Bellieny-Rabelo et al. 2016). *Phaseolus vulgaris:* Common bean gene expression atlas (http://plantgrn.noble.org/PvGEA/) (O’Rourke et al. 2014). *Me. truncatula: Medicago truncatula* Gene Expression Atlas (MtGEA) (Benedito et al. 2008). *Ar. thaliana:* AtGenExpress (Schmid et al. 2005). Rice: Rice Express ion Database; (http://expression.ic4r.org). Maize and *Se. moellendorffii* gene expression data were obtained from recent publications (Stelpflug et al. 2016; Zhu et al. 2017).

